# Attention explores space periodically at the theta frequency

**DOI:** 10.1101/443341

**Authors:** Mehdi Senoussi, James C. Moreland, Niko A. Busch, Laura Dugué

**Author notes:** Corresponding author: Mehdi Senoussi, Current address: Department of Experimental Psychology, Ghent University, H. Dunantlaan 2, 9000 Ghent, Belgium.

## Abstract

Voluntary attention is at the core of a wide variety of cognitive functions. Attention can be oriented to and sustained at a location, or reoriented in space to allow processing at other locations – critical in an ever-changing environment. Numerous studies have investigated attentional orienting in time and space but little is known about the spatio-temporal dynamics of attentional reorienting. Here, we explicitly manipulated attentional reorienting using a cueing procedure in a 2-AFC orientation discrimination task. We interrogated attentional distribution by flashing two probe stimuli with various delays between the pre-cue and target stimuli. Then, we used the probabilities of both probes and none of the probes being correctly reported to solve a second-degree equation, which estimates the report probability at each probe location. We demonstrated that attention reorients periodically at ∼4 Hz (theta) between the two stimulus locations. We further characterized the processing dynamics at each stimulus location, and demonstrated that attention samples each location periodically at ∼11 Hz (alpha). Finally, simulations support our findings and show that this method is sufficiently powered, making it a valuable tool for studying the spatio-temporal dynamics of attention.

## INTRODUCTION

Despite the impression that our visual perception is seamless and continuous across time, mounting evidence suggests that this is an illusion; information is sampled periodically at low frequencies (theta: 4-7 Hz and alpha: 8-12 Hz). Specifically, the alpha and theta rhythms seem to coexist in the brain and support different functions (Dugué, Beck, Marque, & VanRullen, in print; Dugué & VanRullen, 2017; Dugué, Xue, & Carrasco, 2017; VanRullen, 2016). Alpha has been related to sensory aspects of visual perception, and was first described as the natural frequency of the occipital pole (Rosanova et al., 2009). Recent studies further proposed multiple alpha sources (i.e. occipital and parietal) serving distinct functional roles (e.g. (Chaumon & Busch, 2014; Gulbinaite, Viegen, Wieling, Cohen, & VanRullen, 2017; Sokoliuk et al., 2018)). Theta appears to be related to attentional sampling (for review VanRullen, 2016). Specifically, a series of recent research used exploration tasks including visual search (Dugué, Marque, & VanRullen, 2015; Dugué, McLelland, Lajous, & VanRullen, 2015; Dugué & VanRullen, 2014; Dugué, Xue, et al., 2017), priming (Huang, Chen, & Luo, 2015), exogenous (involuntary) spatial attention (Chen, Wang, Wang, Tang, & Zhang, 2017; Landau & Fries, 2012; Landau, Schreyer, van Pelt, & Fries, 2015), endogenous (voluntary) spatial (Song, Meng, Chen, Zhou, & Luo, 2014) and feature-based attention (Fiebelkorn, Pinsk, & Kastner, 2018; Fiebelkorn, Saalmann, & Kastner, 2013; Helfrich et al., 2018) tasks, designed to manipulate covert attention – selective processing of information in the absence of eye movements (Carrasco, 2011, 2014) – and suggested that visual information is sampled periodically at the theta frequency when attentional exploration of the visual space is required (for review see Dugué & VanRullen, 2017).

Recent electrophysiology in macaque monkeys and in humans using ECoG showed that attention-related theta rhythm involves both sensory (V1 and V4; Kienitz et al. (2018); Spyropoulos, Bosman & Fries (2018)) and frontal (Fiebelkorn et al., 2018; Helfrich et al., 2018) cortices. Specifically, Kienitz et al. (2018) propose that theta activity can emerge in sensory areas from competitive receptive field interactions. Fiebelkorn et al. (2018) argue instead that the theta rhythm characterizes functional interactions between the frontal-eye-field and the lateral intraparietal areas. Similarly, we recently proposed that theta rhythmicity may emerge from iterative connections between sensory and attentional regions (Dugué et al., in print; Dugué & VanRullen, 2017).

Together, previous findings suggest that the periodic modulation of attentional processing in time is related to the periodic sampling of visual information in space. Attentional exploration has previously been assessed by manipulating attentional orienting, and then measuring performance using a single stimulus, either at the attended location or at another, unattended location. Here, we intended to clarify whether attention operates at a single location at a time, sampling sequentially across locations, or samples multiple locations simultaneously (VanRullen, 2013). To do so, we directly manipulated attentional reorienting. Reorienting is critical in a dynamic environment that changes rapidly at short timescales, along with observers’ goals and priorities. Characterizing the dynamics of attention’s ability to shift and enhance vision across multiple points of interest is crucial for understanding the limits and capacity of the visual system. Importantly, there is no reason to assume that endogenous orienting (i.e. voluntary engagement of attention on a given spatial location) behaves equally to endogenous reorienting (i.e. first necessitating disengagement of the attention focus from one location to shift and re-engage onto another location; Posner (1988)).

Recently, Dugué et al. (2016) used Transcranial Magnetic Stimulation (TMS) to directly assess the interplay between temporal periodicity and sequential spatial exploration. They investigated the dynamics of endogenous (voluntary), spatial attention. Using a cueing procedure, they explicitly manipulated attentional orienting – attending to one location within a trial – and reorienting – shifting attention from one stimulus location to another, previously unattended location (Corbetta, Patel, & Shulman, 2008; Dugué, Merriam, Heeger, & Carrasco, 2017). By applying TMS at various delays over the occipital pole (V1/V2), they demonstrated that performance in a 2-AFC orientation discrimination task was modulated by occipital TMS periodically at the theta frequency (∼5 Hz) only when attention had to be reallocated from a distractor to a target location. However, due to practical considerations tight to the use of TMS, the frequency resolution was not optimal to allow the precise characterization of the peak frequency of attentional reorienting (eight stimulation delays on a 400-ms window).

Here, we investigated the spatial and temporal dynamics of attentional sampling by explicitly manipulating attentional reorienting. In a psychophysical task, we probed the state of attentional allocation during the course of the trial with high temporal resolution. Two probes were flashed at various delays after the offset of two grating patches. Participants performed a 2-AFC orientation discrimination task on one of the two grating patches, with voluntary attentional orienting and reorienting manipulated using a central cue. Assessing performance on the probes, we used a probability estimation method consisting in solving a second-degree equation using the probabilities of both probes and none of the probes being correctly reported to estimate the amount of attention allocated to the different stimulus locations over time. Critically, by manipulating the probe configuration we were able to analyze both attentional reorienting between stimulus locations, as well as information sampling at each location independently. This approach, first introduced by Dubois, Hamker & VanRullen (2009) and then successfully applied to investigate the spatio-temporal deployment of attention during visual search (Dugué et al., 2015; Dugué, Xue, et al., 2017), allowed us to demonstrate that attentional distribution was periodically modulated over time at the theta frequency (∼4 Hz), only when attention had to be reoriented. By explicitly manipulating the reorienting of endogenous attention, and using a fine-tuned psychophysical and analytical approach, these results suggest that the periodicity in task performance was due to the sequential reorienting of attention from one stimulus location to another, and not the independent sampling of either location. Importantly, our method allowed us to also show that each stimulus location was processed periodically at the alpha frequency (∼11 Hz), suggesting a functional dissociation between the alpha and theta rhythms.

## METHODS

### Observers

Thirteen human observers (M ± SD = 20.9 ± 0.8 years old, range: 20-22 years old; 9 females) were recruited for this experiment, a similar number of participants used in prior studies using the same analysis approach (Dugué et al., 2015; Dugué, Xue, et al., 2017). The decision to stop recruiting new observers was not based on preliminary analyses of the data. We selected a study design in which each participant underwent a large number of trials (1,872 trials total). Due to technical issues during data recording, two observers were excluded from the analysis. All observers had normal or corrected-to-normal vision. They provided written informed consent and received monetary compensation for their participation. All procedures were approved by the CERES (Conseil d’Évaluation Éthique pour les Recherches En Santé) ethics committee of Paris Descartes University. All research was performed in accordance with the relevant guidelines and regulations from the committee.

### Apparatus

Observers sat in a dark room, 57.5 cm from a calibrated and linearized CRT monitor (refresh rate: 85 Hz; resolution: 1280 × 1024 pixels). A chinrest was used to stabilize head position and distance from the screen. Visual stimuli were generated and presented using MATLAB (The MathWorks, Natick, MA) and the MGL toolbox (http://gru.stanford.edu/doku.php/mgl/overview).

### Procedure

Data collection took part across five sessions conducted on five consecutive days. Note that the collected data sets are available through an Open Science Framework repository (https://osf.io/2d9sc/?view_only=6ef3f85d9f944d27b23fc7af5a26f087). Matlab code to replicate the experiment, analyses and figures is provided through GitHub (https://github.com/mehdisenoussi/PECAR). The first session was used to familiarize observers with the experimental protocol. In the four remaining 1-hour sessions, observers performed the main task.

Observers performed a 2-AFC orientation discrimination task as depicted in **Fig1A**. They were instructed to fixate on a white cross at the center of the screen. Eye position was monitored using an infrared video-camera system (EyeLink 1000, SR Research, Ottawa, Canada) to ensure all observers maintained fixation throughout each trial (critical when studying covert attention). Stimulus presentation was contingent upon fixation. Any trials in which observers broke fixation (defined as an eye movement ≥1.5° from the center of the fixation cross or if the observer blinked; M ± S.D. = 15% ± 8%) was canceled and then repeated at the end of each experimental block. Note that eye movements ≤1.5° from the center of the fixation cross (i.e. microsaccades, MS) were not detected online. However, a post-hoc analysis showed that there were inconsequential (M ± S.D. = 0.05 ± 0.01 MS/trial when considering the period between the offset of the grating stimuli and the onset of the probes; see below).

**Figure 1.**
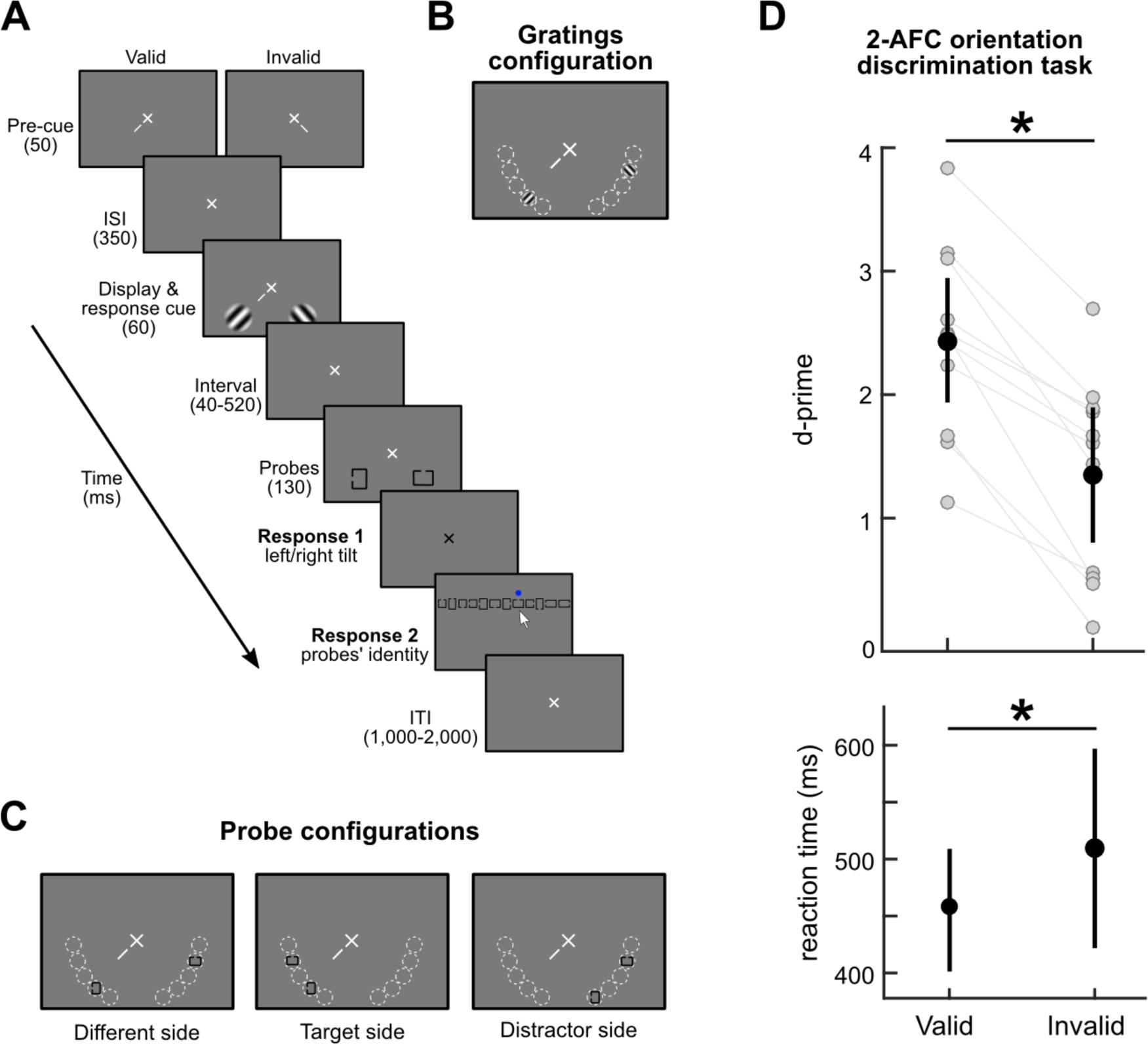
Experimental protocol. (A) Trial sequence: each trial consists of 1) pre-cueing, 2) a 2-AFC orientation discrimination task, and 3) a probe identification task. (B) In each quadrant, the grating appeared in one of five possible positions represented by the dashed white circles (the circles are for illustration purposes only and were not present in the actual experiment). (C) The probes appeared in one of three possible configurations: 1) one probe in each quadrant, or both probes on the same side as 2) the target, or 3) the distractor grating. (D) Performance in the 2-AFC orientation discrimination task. Top panel: d-prime as a function of validity condition (valid and invalid). Each gray dot represents an individual observer. Black dots represent the average across observers. Error bars reflect 95% confidence intervals. D-prime are significantly higher in the valid than invalid condition (t(10)=7.57, p<0.0001, Cohen’s d = 1.368). Bottom panel: median reaction time across observers. Error bars reflect 95% confidence intervals. Reaction times are significantly faster in the valid than invalid condition (t(10)=-2.54, p=0.031, Cohen’s d = 0.479). Results confirm that attention was successfully manipulated, with no speed-accuracy trade-off.

A central endogenous pre-cue presented for 50 ms instructed observers to deploy their covert attention toward the left or right bottom quadrant. The pre-cue was followed by a 350 ms inter-stimulus interval (ISI) – sufficient time to deploy voluntary attention (Busse, Katzner, & Treue, 2008; Carrasco, 2011; Cheal & Lyon, 1991; Dugué, Merriam, et al., 2017; Liu, Stevens, & Carrasco, 2007; Müller & Rabbitt, 1989; Nakayama & Mackeben, 1989; Pestilli, Ling, & Carrasco, 2009). Two stimuli were then presented for 60 ms: sinusoidal gratings windowed by a raised cosine (15% contrast, 3 cycles per degree, at 4° eccentricity, on a gray background). They were tilted randomly clockwise or counter-clockwise relative to the vertical. The tilt angle was chosen for each individual using a staircase procedure performed during the first session using a neutral pre-cue (two pre-cues: one pointing to the right and the other one to the left quadrant, half the size of the single pre-cue) to achieve ∼75% accuracy (averaged tilt angle, M ± S.D = 7.02 ± 4.02 degrees) independently for all observers. The target grating, i.e. the stimulus for which observers had to report the orientation, was indicated by a central response cue pointing either toward the left or the right bottom quadrant. Two gratings were always presented, one on each quadrant (in each quadrant the location was randomly selected from one of five possible locations, as depicted in **Fig1B**). A trial was valid when the quadrant in which the target grating appeared matched the pre-cued quadrant (75% of the trials). A trial was invalid when the target appeared at the uncued quadrant (25% of the trials), requiring reorienting attention endogenously to the opposite quadrant. For each participant there were a total of 1404 valid and 468 invalid trials. Note that for all analyses we used all available trials. However, to ensure that any observable difference between the valid and invalid trials conditions is not due to different amount of trials, we also performed the main analysis (**Fig2**, left column) by sub-sampling for each participant the amount of valid trials to match it with the amount of invalid trials –i.e. for the valid trial condition only, we re-ran the analysis on 468 randomly selected valid trials (1,000 iterations)– and found similar results.

**Figure 2.**
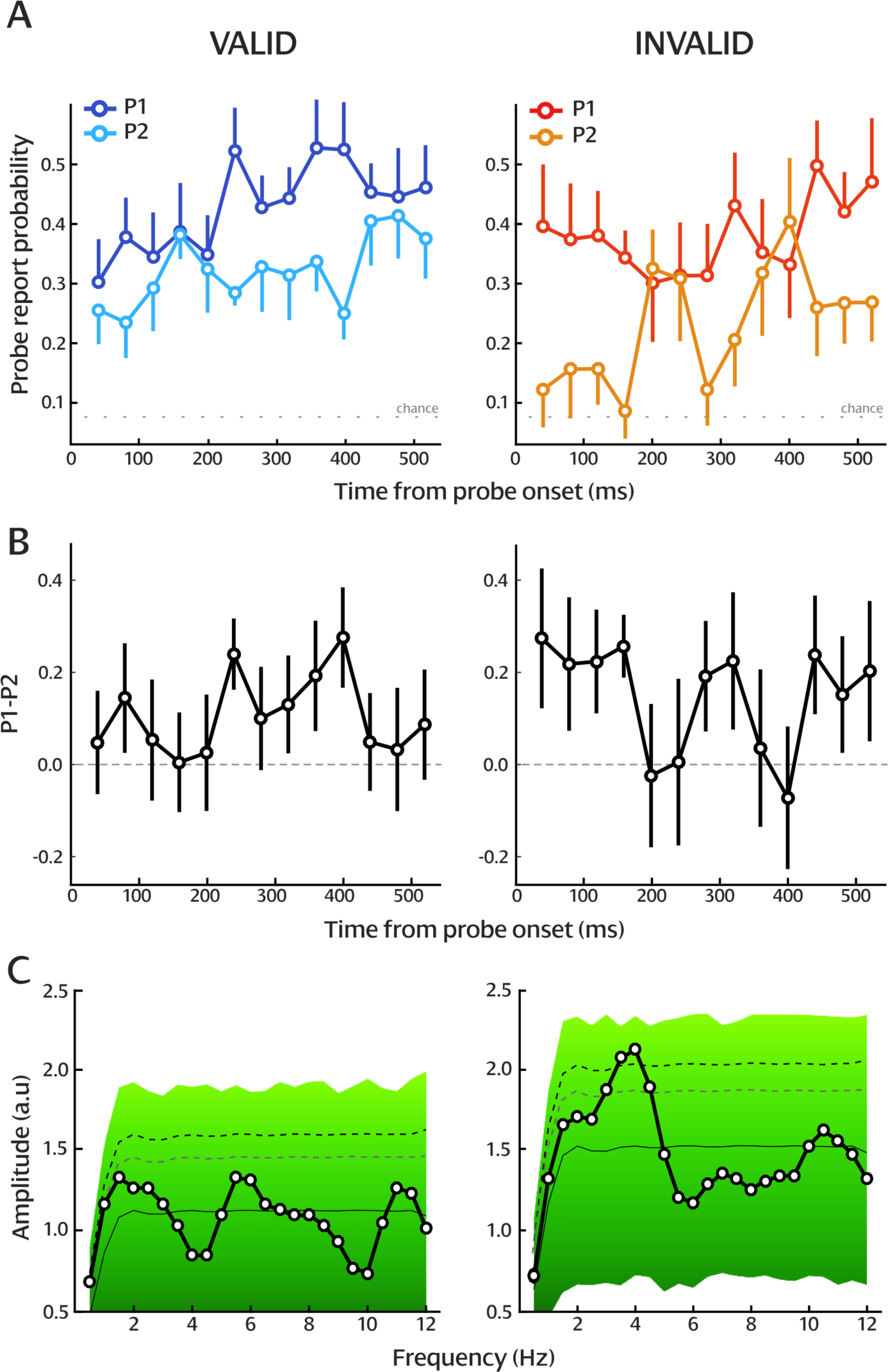
Dynamics of attentional orienting and reorienting. (A) P1 and P2 by delays for valid (left column) and invalid (right column) conditions (probability estimation from the probe task). The dotted line represents the probability of reporting correctly the probe by chance (0.083). Error bars reflect 95% confidence intervals. Only one half of the error bars is shown to reduce visual clutter. (B) Difference between P1 and P2. The dashed line represents a zero difference. Error bars reflect 95% confidence intervals. (C) Amplitude spectrum of the P1-P2 difference and results of the 100,000 surrogate analysis for valid and invalid conditions. In each graph, the solid line represents the average of the surrogate analysis. The black-to-green background represents the distribution of surrogate values, i.e. the level of significance of each frequency component. The gray dashed line represents p < 0.05. The black dashed line represents p < 0.05 after Bonferroni correction.

Two Landolt C’s, squares or rectangles with an aperture on one of the sides (12 possible probes: four squares, four horizontal rectangles and four vertical rectangles), were presented for 130 ms after one of 13 possible delays after the onset of the orientation discrimination display (ISI: 40-520 ms, in 40 ms increments; 144 trials per delay). Upon offset of the probes, the fixation-cross turned black to indicate the beginning of the response window during which observers had to report the tilt orientation of the target (clockwise or counter-clockwise to vertical). Auditory, positive (high tone) or negative (low tone), feedback was provided to the observers. We calculated d-prime to measure accuracy as our main dependent variable. To rule out any speed-accuracy trade-offs, we also report median reaction time (RT) from the onset of the response window as secondary dependent variable. The fixation-cross disappeared after the observers responded and a horizontally aligned array of the 12 possible probes appeared. The order of probes’ identity in the array was randomized between trials. Observers used the computer mouse to report probes’ identity by clicking on each of the two probes they perceived (see **Fig1A**). When observers clicked on a given probe in the array a blue dot appeared for 500 ms above the probe to indicate that it was selected. It was possible for the two probes to be identical (observers were aware of this possibility), in which case observers had to click twice on the same probe. There was no feedback for performance on the probe task in order to ensure that observers prioritized the first task. For each trial, we recorded whether the observer reported none, one, or both probes correctly.

There were three probe location conditions: 1) the two probes were presented in opposite quadrants (randomly located in one of the 5 possible locations within the quadrant, **Fig1C** left panel), or both in the same quadrant as 2) the target (**Fig1C** middle panel) or 3) the distractor (opposite side, **Fig1C** right panel). The analysis performed on trials in which one probe was presented in each quadrant assayed attentional reorienting, while the analysis performed on the other two conditions informed information sampling at each location. All analyses were performed on trials in which observers correctly responded in the 2-AFC task (grating’s orientation) to ensure that they followed the primary task’s instructions, and that attention was successfully deployed to the target location. Crucially, the Landolt C’s were used to probe the state of attentional distribution (see **Probability estimates** section). Specifically, this manipulation allows interrogation of the spatial location of the attention focus at various delays over the course of the trial. Note that our approach allowed us to determine whether both probes received the same amount of attention, or whether one received more than the other one on average across all trials. Yet, we remain agnostic of the exact location of the attentional focus on a trial-by-trial basis.

### Probability estimates

We aimed to investigate the spatio-temporal behavior of attention during a task in which we explicitly manipulated the orienting and reorienting of endogenous attention. Specifically, performance to identify the two probes presented at different locations shortly after the gratings informs us on how attention was differentially distributed in the visual field. In other words, we ask whether attention is sampling both probe locations equally at each time point, or whether the distribution of attentional resources between probe locations is modulated over time, suggesting that attention alternates between probe locations in time.

To examine this question, we used the method previously introduced by Dubois et al. (2009) and then successfully applied in visual search tasks by Dugué and colleagues (2015; Dugué, Xue, et al., 2017) to characterize attentional deployment. This probability estimation method allows extracting an estimate of attentional distribution when the location of attention is unknown. Note that in our experiment we have two types of probe configurations: either both probes were presented within the same quadrant, or in opposite quadrants. Our paradigm allows access to the relative attentional distribution between quadrants over time (cued vs. uncued quadrant), but not within quadrant (five possible stimulus positions). We thus employed a probability estimation method able to estimate attentional distribution for all probe configurations.

The probabilities of attending to each of the probed locations, called P1 (probability of reporting the probe at the most attended location) and P2 (probability of reporting the probe at the least attended location), were determined using the probabilities of reporting both probes correctly (Pboth) and none of the probes correctly (Pnone).

The probability of getting both probes correct is,

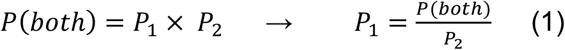

Note that the relation of independence between the probes is ensured by the task allowing them to be identical (i.e. the choice of the second probe does not depend on the choice of the first one). The probability of getting neither correct is,

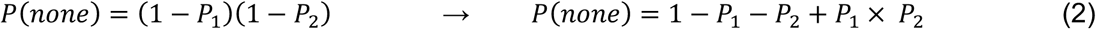

Substituting (1) into (2):

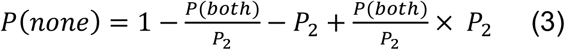

Moving all to one side of the equation:

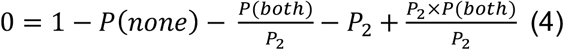

Multiplying all by –*P*_2_:

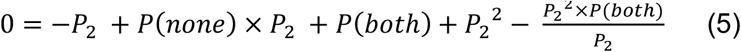

And rearranging:

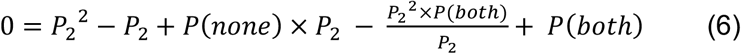

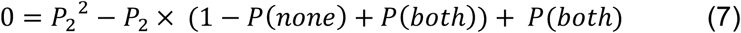

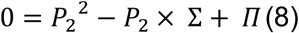

With:

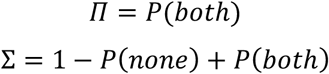

Equation 8 is now quadratic and can be solved using the quadratic formula for *P*_2_. Because *P*_1_and *P*_2_ are symmetric in the equations above, the two solutions of (8) are *P*_1_and *P*_2_.

Calculating the discriminant (Δ) as follow will allow us solving Equation (8)

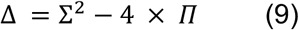

And the solutions of equation (8) are:

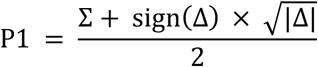

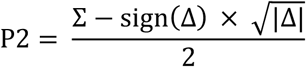

Note that due to noise in responses and performance, and a relatively low number of trials per condition, negative values of Δ were sometimes obtained, which can produce non-real solutions. To address this problem, we followed the same approach than Dugué and colleagues (2015; 2017) and used |Δ| and sign(Δ) thus allowing P1 and P2 values to be negative.

### Evaluating the mathematical approach to calculate attentional distribution

For comparison purposes with the results of the previous studies, we present the results as a function of P1 and P2 using the absolute value and the sign of the discriminant (see previous section). As developed in the previous studies, this is a more conservative solution than assigning a zero value to Δ or taking its absolute, which would artificially increase differences between P1 and P2. This approach also has the advantage of giving a straightforward, intuitive interpretation, i.e. if P1 equals P2 then both probe locations received the same amount of attention; however, if P1 greater than P2, then one location received more attention than the other one.

Yet, to further test our probability estimates we also performed the analysis of the discriminant. As shown below, the discriminant is the square of the difference between *P*_1_and *P*_2_ in the case where coefficient a = 1, and is an unbiased estimate.

Beginning with the quadratic formula,

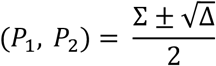

We can subtract the negative solution from the positive to describe *P*_1_ – *P*_2_,

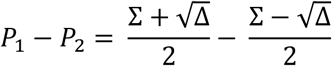

Which reduces to,

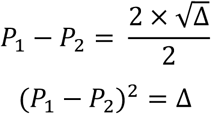

This means that the discriminant as it was used in the analyses of the previous papers is equal to (*P*_1_ – *P*_2_) before the absolute value was taken. Then, if there is no difference between the probability of reporting either probe, then *P*_1_ – *P*_2_ = 0, and so will the discriminant, (*P*_1_ – *P*_2_)^2^= 0. If performance on one probe depends on the other, and correctly reporting one makes reporting the other correctly less likely, then the discriminant will scale (non-linearly) with the size of this difference.

### Spectral analysis

To analyze the temporal dynamics of attentional distribution in valid and invalid conditions, we computed a Fast Fourier Transform (FFT –we decomposed the behavioral data from the time domain into frequency components to estimate an amplitude spectrum, i.e. the amount of each frequency present in the original data (e.g. (Chota et al., 2018; Dugué et al., 2015, 2016; Dugué, Xue, et al., 2017; Fiebelkorn et al., 2013; Ho, Leung, Burr, Alais, & Morrone, 2017; Landau & Fries, 2012; Song et al., 2014; Zhang, Morrone, & Alais, 2019)), for each observer, on the difference between P1 and P2, and then averaged the resulting amplitude spectra across observers. Note that, here, we did not analyze the absolute values of P1 and P2. Instead, we were interested about the difference between P1 and P2, which represents the relative distribution of attention across the two probes regardless of their actual location, i.e. a difference of zero indicates that both probe locations receive the same amount of attentional resources (uniform attentional distribution) while a positive difference indicates that one probe location receives more resources than the other one (non-uniform attentional distribution).

The data of each observer was average-padded to increase the frequency resolution, i.e. values corresponding to the average of the P1-P2 difference across delays were added on either side of the empirical data points. Specifically, the 13 time points, spanning 480 ms, were padded to get a 2000 ms segment, thus adding 18 data points before the first data point and 19 data points after the last one. Note that we also performed the analysis on non-padded data and obtained similar results.

To calculate statistical significance of each frequency component, we used permutation tests (100,000 surrogates of the index for each participant), under the null hypothesis of a random temporal structure of the P1-P2 difference. We shuffled the delays, padded the data as described above, and computed the FFT over each surrogate. The amplitude of the surrogate FFT results was then sorted in ascending order. We used a statistical threshold at p < 0.05, corrected for multiple comparisons (6 frequency components corresponding to the true frequency resolution of the P1-P2 difference) using Bonferroni correction (833^rd^ highest value, in other words p < 0.00833).

Finally, in addition of the probability estimation method that we use to investigate the temporal dynamics of attentional distribution within and between quadrants, we also analyzed the temporal dynamics of probe report accuracy for trials in which one probe appeared in each quadrant (**Fig2C** left panel). For these trials, we computed the FFT for each observer on the difference between probe report accuracy in the attended quadrant (indicated by the pre-cue) and in the unattended quadrant, and then averaged the resulting amplitude spectra across observers. We again padded to the average, and significance was calculated by shuffling the probe report accuracy difference across delays.

## RESULTS

### Grating task

We first evaluated the performance in the 2-AFC orientation discrimination task (**Fig1D**) in order to ensure that observers correctly performed the task and that attention was successfully manipulated. We computed d-prime for reporting the tilt orientation of the target grating in the valid and invalid conditions and observed significantly higher sensitivities for the valid than invalid condition (two-tailed, paired-sample t-test, *t*(10) = 7.57, *p* < 10^−4^, 95%-CI, confidence interval = [0.76 1.4], Cohen’s d = 1.368). There was also a significant difference in median reaction times between valid and invalid condition, i.e. observers responded faster in the valid condition (*t*(10) = −2.54, *p* = 0.031, 95%-CI = [−0.10 −0.006], Cohen’s d = 0.479). Both variables reflected an attentional facilitation in the 2-AFC orientation discrimination task demonstrating that attention was successfully manipulated, with no speed-accuracy trade-offs.

### Probe task

We measured performance in reporting both probes (Pboth) and neither of the probes (Pnone) correctly. To investigate the dynamics of attentional reorienting, we used Pboth and Pnone to estimate probe report probabilities for the condition in which the probes were presented in opposite quadrants (see **Methods** section). We observed that P1 and P2 were marginally significantly different in the valid condition (two-way repeated-measures ANOVA: F(1,10) = 4.405, p = 0.062, η^2^ = 0.30; **Fig2A**, left panel). This difference was significant in the invalid condition (F(1,10) = 22.976, p = 0.0007, η^2^ = 0.69; **Fig2A**, right panel). Both were well above chance (chance in probe report probability = 0.083). We argue that attention in the probe task was non-uniformly distributed across the two stimulus locations.

To assess the temporal dynamics of attentional distribution, we calculated the difference between P1 and P2 (**Fig2B**), separately for valid and invalid trials. We then performed an FFT on the P1-P2 difference for each observer, and analyzed the resulting averaged amplitude spectra (**Fig2C**; see **Methods** section). In the valid condition, there was no significant peak frequency. In the invalid condition, there was a significant peak at 3.5 and 4 Hz (p < 0.05, Bonferroni corrected) suggesting that the P1-P2 difference of each observer was modulated periodically at the theta frequency.

To further validate our results, we compared valid and invalid conditions (**Fig3A**) for the P1-P2 difference and replicated it on the discriminant (see **Methods** section). For the P1-P2 difference, the amplitude of the 4 Hz component for each observer was significantly higher in the invalid than valid condition (one-tailed, paired-sample t-test, t(10) = 3.86, p = 0.001, 95%-CI = [0.5296 1.977], Cohen’s d = 1.647), consistent with the results of **Fig2C**. For the discriminant, the amplitude of the 4 Hz component was also significantly higher for the invalid than the valid condition (**Fig3B**; one-tailed, paired-sample t-test, *t*(10) = 3.36, *p* = 0.004, 95%-CI = [0.2691 1.3298], Cohen’s d = 1.383), confirming the validity of our method.

**Figure 3.**
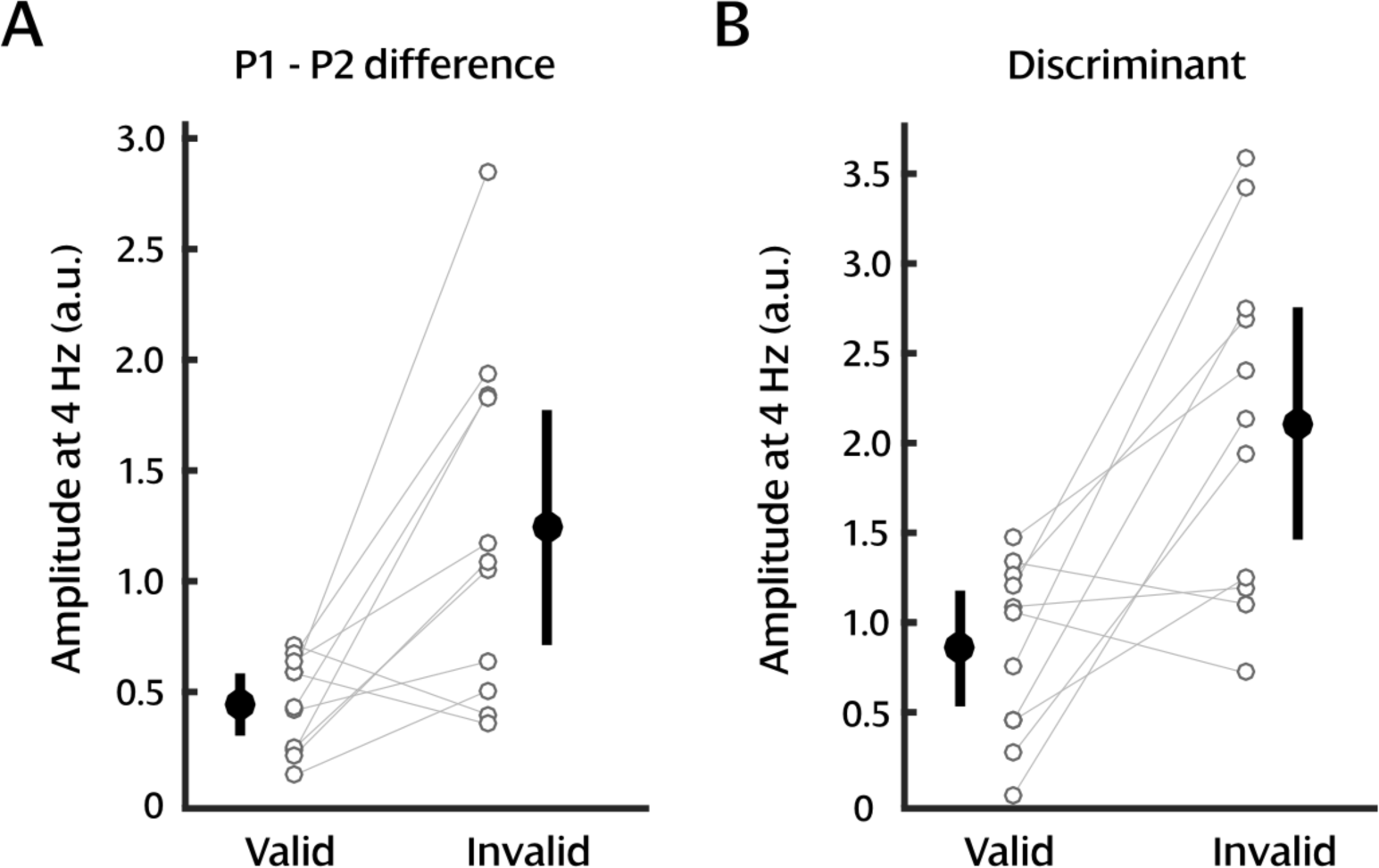
Analysis comparison. An FFT was performed for each observer separately for both the P1-P2 difference (A) and the Discriminant (B), for valid and invalid conditions. Empty black circles represent the amplitude of the 4 Hz component for individual observers. Black filled circles represent the average across observers. Error bars reflect 95% confidence intervals. The difference between valid and invalid conditions was significant for both the P1-P2 difference (t(10) = 3.86, p = 0.001, Cohen’s d = 1.647) and the Discriminant (t(10) = 3.36, p = 0.004, Cohen’s d = 1.383).

To assess the temporal dynamics of attentional distribution, we estimated probe report probability at the most (P1) and least (P2) attended location using the analysis approach described in the **Methods** section, which is agnostic to the actual probe location. However, because of our cueing manipulation, we know which probe is presented in the attended quadrant (i.e. quadrant indicated by the pre-cue) and which one is presented in the unattended quadrant. To confirm the results from the previous analysis, we thus performed the same spectral decomposition directly on the difference in probe report accuracy between the attended and unattended quadrant, taking again only trials in which the probes appear in opposite quadrants (**Fig4**; see **Methods** section). In the valid condition, there was no significant peak in frequency (**Fig4A**). In the invalid condition, there was a significant peak at 3 Hz (**Fig4B**; p = 0.022; although it does not withstand Bonferroni correction), in line with the periodic modulation previously observed in the P1-P2 difference (3.5 - 4 Hz; **Fig2C**).

**Figure 4.**
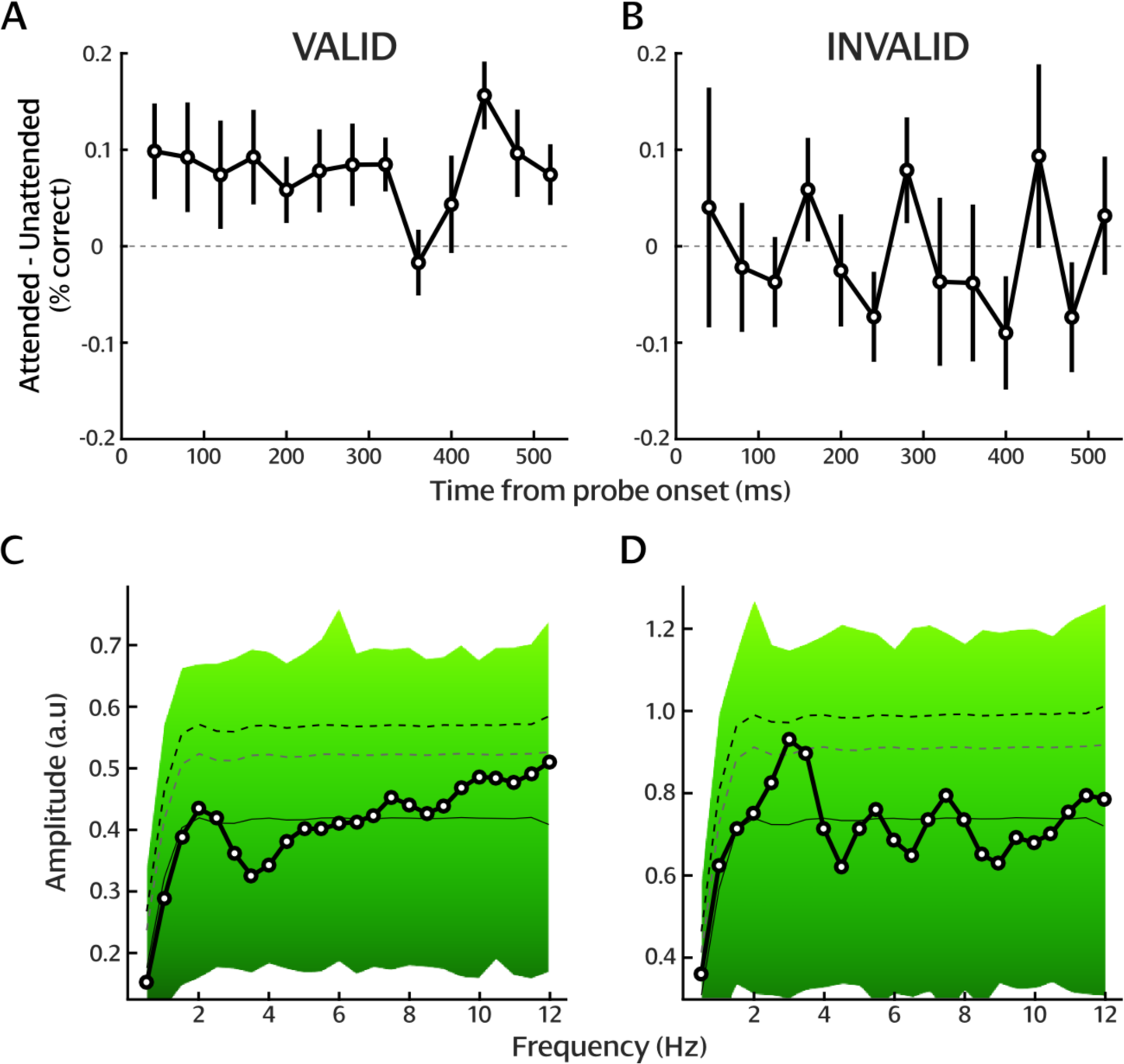
Dynamics of attentional distribution measured using probes report accuracies. Difference between attended (indicated by the pre-cue) and unattended probe report accuracies for (A) valid and (B) invalid trials. The dashed line represents a zero difference. Error bars reflect standard error to the mean across observers. Amplitude spectra of the difference in probe report accuracy between trials in which the probe appeared in the attended and unattended quadrant, for (C) valid and (D) invalid trials, and results of the 100,000 surrogates analysis. The solid line represents the average of the surrogate analysis. The black-to-green background represents the distribution of surrogate values, i.e. the level of significance of each frequency component. The gray dashed line represents p < 0.05. The black dashed line represents p < 0.05 after Bonferroni correction.

Note that this effect is not as strong as the one obtained using the probability estimation method. As described in **Fig1C**, the probe could appear at five possible positions in the attended or unattended quadrant, and potentially at a different location from the grating in the quadrant. For both analyses, we combine all trials together, regardless of the actual location of the probe within each quadrant. The probability estimation method is insensitive to specific probe location, i.e. it estimates the relative attentional distribution across spatial locations. However, when examining probe reportmaccuracies, we estimate the absolute probability of reporting a single probe, which is arguably influenced by the specific stimulus configuration.

Critically, to investigate attentional distribution within each quadrant, we analyzed trials in which both probes were presented in the same quadrant as the target or the distractor (**Fig1C** middle and right panels). In this case, analyzing probe report accuracies cannot inform the attentional distribution within the quadrant since we do not know where attention is allocated within the quadrant –an entire quadrant was pre-cued on each trial. Thus, we need to use the probability estimation method. Specifically, we performed the same analysis as in **Fig2C** for trials in which both probes were presented in the same quadrant, i.e. same side as the target or the distractor. Note that, here, we used the probability estimation method to evaluate, in each quadrant, whether one probe is processed more efficiently than the other one, independently of our manipulation of attentional distribution. The results show a significant effect in the valid trial condition at low frequencies (<1.5 Hz; p<0.0234; although it does not withstand the Bonferroni correction; **Fig5A**). This effect presumably corresponds to a slow trend in the P1-P2 difference over the course of the trial, and not a true periodic modulation. In the invalid trial condition, we observe a significant periodic modulation of the P1-P2 difference in the alpha frequency range (10-11 Hz; p=0.03; although it does not withstand the Bonferroni correction; **Fig5B**). We finally show that this periodic modulation at the alpha frequency is present for both 1) trials in which the probe appears in the same quadrant as the target (10 Hz; p=0.0110; although does not withstand the Bonferroni correction; **Fig5C** top panel), and 2) trials in which the probe appears in the same quadrant as the distractor (11 Hz; p=0.0074; **Fig5C** bottom panel).

**Figure 5.**
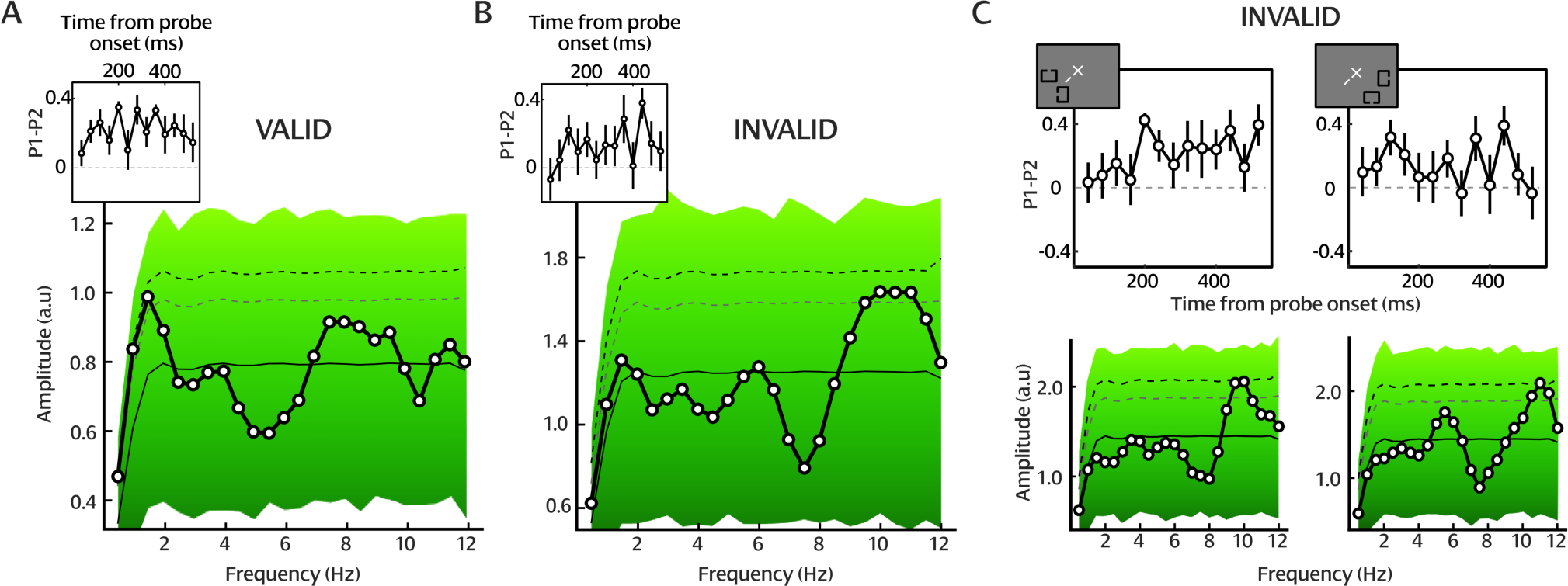
Dynamics of information processing in each visual quadrant. P1-P2 difference, corresponding amplitude spectra, and results of the 100,000 surrogate analysis for (A) valid, (B) invalid trial conditions. The same analysis is presented for the invalid trial condition separately for trials in which both probes are presented on the target side (C left), and trials in which both probes are presented on the distractor side (C right). In each graph of the P1-P2 difference, the dashed line represents a zero difference, and error bars reflect 95% confidence intervals. In each the graphs representing the amplitude spectra, the solid line represents the average of the surrogate analysis. The black-to-green background represents the distribution of surrogate values, i.e. the level of significance of each frequency component. The gray dashed line represents p < 0.05, the black dashed line represents p < 0.05 after Bonferroni correction.

### Power analysis

The sequential attentional reorienting interpretation of the results suggests that within a trial the probability of detecting the probe’s identity is dependent across locations, i.e. that if the observers correctly identify a probe at one location, they are less likely to identify the probe at another. This dependency is lost if the probe responses to one location are randomly permuted. The result is the same overall percent correct for each probe but probe responses within each trial are independent.

To evaluate the statistical power of the probability estimation method, we tested the null hypothesis of no dependency between the two probe responses within a trial. We generated a surrogate distribution under this null hypothesis by repeatedly (10,000 repetitions) permuting the probe responses at one location and performing the quadratic analysis on the resulting pairs of probe responses. This null distribution was then compared to a distribution generated by resampling with replacement trial responses while preserving any existing dependency within a trial (**Fig6**). The achieved power is the proportion of sampling repetitions from the resampling P1-P2 difference (**Fig6A** and **6B**) and discriminant (**Fig6C** and **6D**) distributions that are greater than the 95^th^ percentile of the permuted distributions. For the P1-P2 difference the achieved power in the valid condition was 99.4%, and in the invalid condition was 10.5%. For the discriminant the achieved power in the valid condition was 100%, and in the invalid condition was 34.4%. Interestingly, the permuted data does not have a discriminant distribution centered on zero as would be expected if there were no relation between the two probe responses. A possible explanation is that observers had a bias in performance for the first or second probe responses, which is a relation preserved in the permutation method described above.

**Figure 6.**
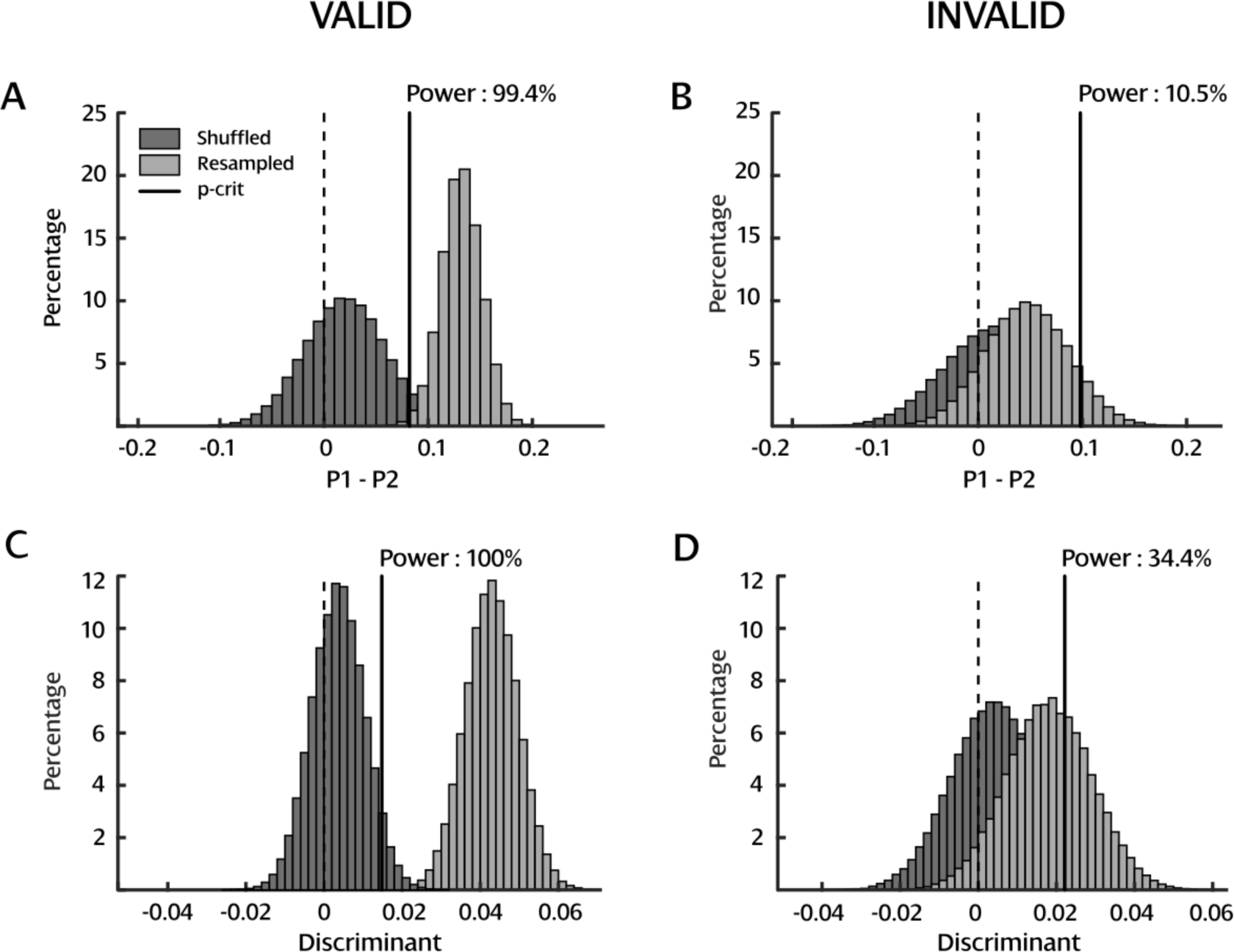
Power analysis. Distributions of P1-P2 difference and discriminant for resampled and permuted trials, for valid (left panels) and invalid (right panels) conditions. Solid vertical lines in each panel indicate the 95^th^ percentile of the permuted distribution. Proportion of resampled repetitions to the right of the solid line indicates the power achieved.

In conclusion, using this power analysis, we addressed the following question: Is the quadratic analysis approach sufficiently powered to detect a difference between the probability of correctly reporting probe one and probe two? Based on the permutation tests (**Fig6**), we were able to conclude that this method is appropriate and sufficiently powered to detect such difference. This assessment adds support to the literature already published (Dubois et al., 2013; Dugué et al., 2015, 2016).

## DISCUSSION

We investigated the spatio-temporal dynamics of attentional orienting and reorienting using a psychophysical approach with high temporal resolution. By explicitly manipulating covert spatial, endogenous attention, we demonstrated that performance in the invalid condition, when attention needed to be reoriented from the distractor to the target location, was periodically modulated at the theta frequency (∼4 Hz; **Fig2** and **Fig4**). Additionally, we found that stimuli presented in the same quadrant were sampled periodically at the alpha frequency (∼11 Hz; **Fig5**; see **Fig7**). Finally, we performed our analysis on another index of attentional distribution –the discriminant (**Fig3**)– and replicated the effect.

**Figure 7.**
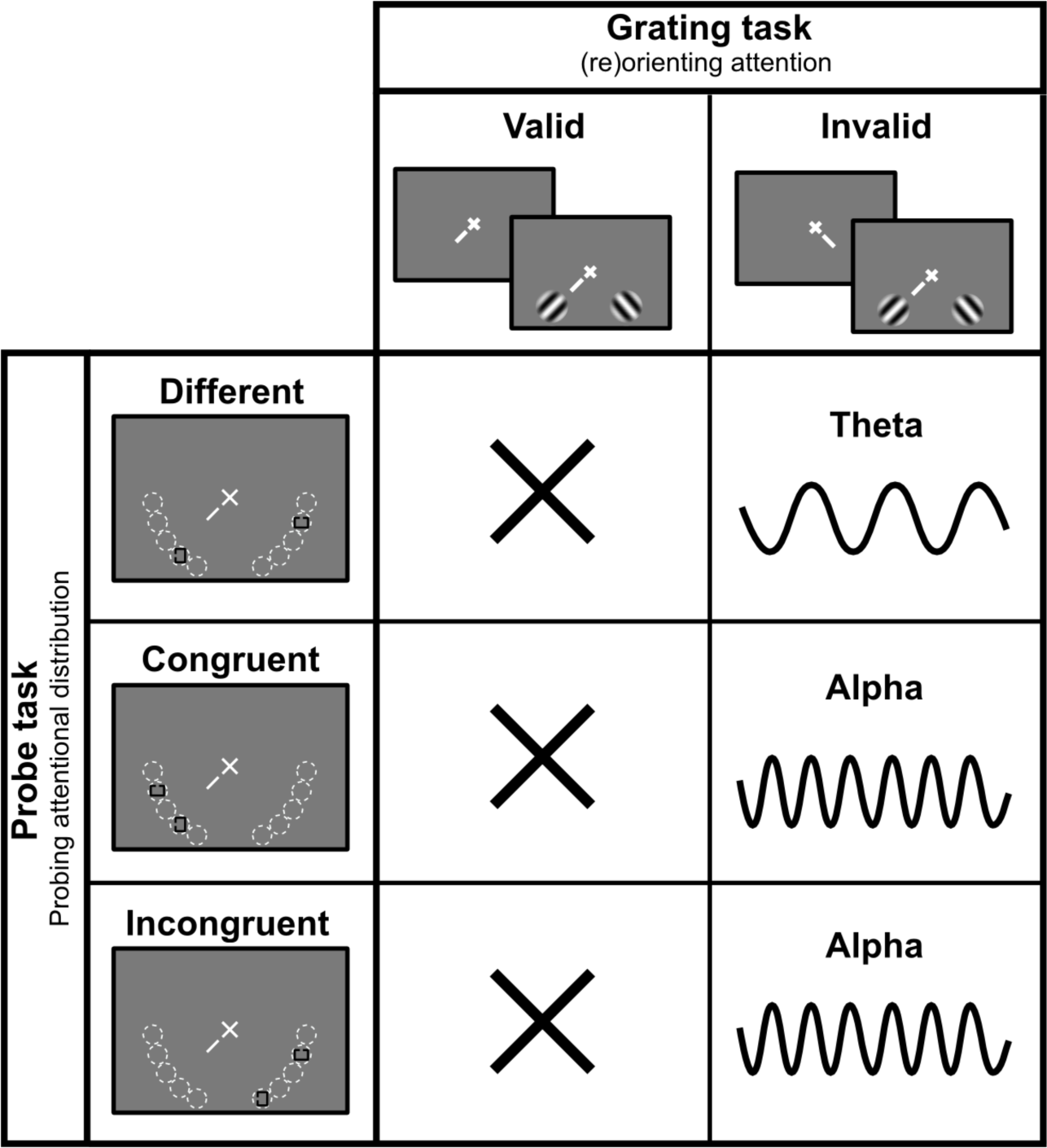
Summary. Attention reorients (invalid, different) periodically between the attended and the unattended location at the theta frequency (∼4 Hz), and each stimulus location (invalid, congruent and incongruent) is sampled periodically at the alpha frequency (∼11 Hz).

Interestingly, our probability estimation approach not only allowed us to investigate the rhythm of attentional reorienting, but also the rhythm of information sampling at each location. We found a functional dissociation between the theta rhythm, observed when analyzing trials in which the probes are presented in opposite quadrants, and the alpha rhythm, observed when analyzing trials in which both probes are presented in the same quadrant. Specifically, our results suggest that the alpha and theta rhythms jointly allow periodic sampling of the visual environment. The ongoing alpha rhythm, i.e. sensory rhythm (∼11 Hz; whose phase may not be reset by stimulus onset, or only partially), would coexist with the theta attentional rhythm (∼4 Hz), and either be recordable concurrently or be masked depending on task relevance. This is consistent with recent literature reviews suggesting that theta corresponds to the rhythm of attentional exploration while alpha rather reflects an ongoing, sensory sampling rhythm (Davidson, Alais, Boxtel, & Tsuchiya, 2018; Dugué & VanRullen, 2017; VanRullen, 2016).

Recent studies have shown that interactions between frequency bands was critical for the brain to orchestrate different aspects of visual and attentional processing (Helfrich, Huang, Wilson, & Knight, 2017; Klimesch, 2018). Specifically, Helfrich et al. (2017) recently observed that phase-amplitude cross-frequency coupling between 2-4 Hz and 8-12 Hz oscillations, i.e. the phase of the 2-4 Hz oscillations modulates the amplitude of the 8-12 Hz oscillations, alters behavioral performance in a visual detection task. Here, because theta and alpha behavioral rhythms were revealed in different conditions, we were not able to test for a potential interaction, and further research will be necessary to investigate this question.

Previous studies have demonstrated the role of theta oscillations in the attentional exploration of the visual space (Busch & VanRullen, 2010; Dugué et al., 2015; Jia, Liu, Fang, & Luo, 2017; Landau et al., 2015; Rollenhagen & Olson, 2005), consistent with the theta periodicity observed here in the invalid condition. These authors have suggested that the observed behavioral periodicity is due to an intrinsic property of the system, i.e. brain oscillations at the theta frequency modulate behavioral performance periodically at the same frequency. A recent psychophysics study explicitly manipulated spatial attentional orienting to assess the behavioral periodicity of attentional sampling with a discrimination task (Song et al., 2014). Such tasks allow assessment of whether the effect of attentional cueing reflects changes in sensory (d’) rather than decisional (criterion) processing. On the contrary, in detection tasks, the challenge is to disentangle whether higher performance is due to facilitation of information coding at that location, probability matching, or a decision mechanism (see Dugué, Merriam, et al., (2017)). The results of the Song et al. study suggest that attention samples information periodically at low frequencies (alpha and theta). Unfortunately, the main dependent variable was reaction time, which can reflect perceptual processing speed, motor anticipation (Correa, Triviño, Pérez-Dueñas, Acosta, & Lupiáñez, 2010), and criterion (Carrasco & McElree, 2001). Here, we explicitly manipulated the reorienting of attention to directly address the following question: is the periodicity in behavioral performance due to the sequential sampling by attention of the different stimulus locations or to the independent sampling of each location? We measured d-prime as main dependent variable and verified that attention in the 2-AFC orientation discrimination task was successfully manipulated for each participant, with no speed-accuracy trade-off.

Critically, we flashed two stimuli at various delays after the 2-AFC task to probe the state of attentional allocation during the course of the trial. Using this manipulation we argue that, in the invalid condition, the response cue (i.e. the reorienting signal) did not reorient attention only once but multiple times, attention periodically switching from one spatial location to another. This surprising result was recently reported in the study by Dugué et al. (2016), which employed a similar approach. They used TMS to interfere with the processing of the target and the distractor stimuli independently, at various delays during the same 2-AFC orientation discrimination task, in order to perturb the orienting or reorienting of attention. They similarly observed that endogenous attention periodically reoriented from the distractor location to the target location (invalid condition) at the theta frequency (∼5 Hz). Here, using a psychophysics approach instead of directly interfering with neural processing, we replicated this finding. Our results point to a slightly lower frequency of reorienting, i.e. ∼4 Hz. This discrepancy might be due to a different temporal resolution between the two studies (i.e. 13 different delays were sampled here, reaching a higher temporal resolution). Taken together, these results suggest that, in this specific 2-AFC orientation discrimination task, attention is not independently sampling each location periodically, but is sequentially sampling the two stimulus positions, alternating from one position to another at the theta frequency.

Interestingly, the specific frequency of attentional exploration, although always in the theta frequency range, varies across studies between 4 and 8 Hz (Busch & VanRullen, 2010; Dugué et al., 2015; Dugué, Xue, et al., 2017; Fiebelkorn et al., 2013; Huang et al., 2015; Landau & Fries, 2012; Landau et al., 2015; Song et al., 2014; van Diepen, Miller, Mazaheri, & Geng, 2016; VanRullen, 2013; VanRullen, Carlson, & Cavanagh, 2007). It is possible that the differences in task parameters could modulate the frequency of the attentional sampling. Recent studies show that the peak frequency of alpha oscillations can be shifted depending on task demands or general cognitive load (Haegens, Cousijn, Wallis, Harrison, & Nobre, 2014; Mierau, Klimesch, & Lefebvre, 2017; Wutz, Melcher, & Samaha, 2018). These studies suggest that a top-down modulation of the alpha peak frequency in occipital areas optimizes sensory sampling and inter-areal communication (see review and model by Mierau et al., (2017)). Although these studies do not explicitly suggest a general mechanism that would also be present in other frequency bands, e.g. theta, it may be the case that modulation in the theta peak frequency due to task demands and top-down factors could explain differences in the precise peak frequency across studies and tasks.

We found a periodic modulation in the invalid but not in the valid condition, i.e. when attention was reoriented but not when it was sustained at the initially cued location, consistent with Dugué et al. (2016). Other studies showed modulations in the probability of detecting a target in conditions where attention was sustained (Fiebelkorn et al., 2013; Rollenhagen & Olson, 2005; VanRullen et al., 2007), although different types of attention were manipulated. As argued in Dugué et al. (2016), it is possible that sampling in the valid condition may have been similarly modulated, but with its own spontaneous phase. In this condition, the orienting process would not have reset the underlying oscillation, and the probes would not bear a phase relation with attention. However, in the invalid condition, the attentional reorienting process would have reset the phase of the oscillation, which could then be revealed by the two probe stimuli. This result challenges the idea that theta oscillations serve as a general attentional rhythm across spatial locations, suggesting that attentional reorienting is crucial to trigger the theta sampling process.

Recently, several studies have investigated the role of microsaccades in brain and behavioral rhythms (Bellet, Chen, & Hafed, 2017; Bosman, Womelsdorf, Desimone, & Fries, 2009; Deouell, 2016; Helfrich, 2017). Specifically, Bellet et al. (2017) recently demonstrated that both the onset time and the direction of microsaccades modulate alpha and beta periodic modulations of reaction times in a detection task. Lowet et al. (2016) further showed that gamma oscillations in areas V1 and V2 of the monkey brain covaried with a microsaccade-related theta oscillation. Together, these findings suggest that microsaccades may organize visual processing through phase reset of brain oscillations (Deouell, 2016; Helfrich, 2017). Our paradigm did not allow for investigating microsaccades (i.e. very low number of microsaccades between the offset of the grating stimuli and the onset of the probes, which is where attentional reorienting takes place). Further studies will thus be needed to investigate the link between microsaccades and the observed theta and alpha behavioral rhythms.

## CONCLUSIONS

Using a probability estimation approach, we not only show that attentional reorienting samples space periodically at the theta frequency (∼4 Hz), but also that each spatial location is sampled periodically at the alpha frequency (∼11 Hz) by an ongoing sensory rhythm. Together, these results support the idea that our brain samples information periodically, at low frequencies. Finally, our analysis approach using probability estimation of stimulus identification to probe the state of the attention system is a valuable, sufficiently powered tool to investigate the spatio-temporal dynamics of information processing.

## ACKNOWLEDGEMENTS

This work was supported by an ANR-DFG grant to L.D. (J18P08ANR00) and N.A.B. (BU 2400/8-1). The authors thank Ian Donovan for his useful comments on the manuscript. The authors declare no competing interests.

